# Stock mislabelling and its impact on phylogenomic analysis in the *Drosophila nasuta* subgroup/species complex

**DOI:** 10.1101/2019.12.30.891523

**Authors:** Peter J. Waddell, Jeremy Wang, Corbin D. Jones

**Affiliations:** School of Fundamental Sciences, Massey University, Palmerston North, New Zealand; Department of Genetics, University of North Carolina, Chapel Hill, North Carolina; Department of Biological Sciences, University of North Carolina, Chapel Hill, North Carolina

## Abstract

In the course of our 28 de novo genome study of the *nasuta*-subgroup of the *immigrans* species group, we have come to suspect that several stocks acquired from the major stock centers may have either been mislabeled or mixed up in other ways. This is a short note to indicate which stocks these are, how they were diagnosed, and how an alignment and assembly free (AFAF) phylogenomic analysis sheds further light on what probably occurred.

## Introduction

While flies of this subgroup are quite distinct from their closest relatives such as the *immigrans* subgroup, *hypocausta* subgroup and *neohypocausta* subgroup, identifying the specific species in the *nasuta*-subgroup faces several challenges, such as cryptic species. However, on the frons, there are three main patterns of pollinosity or shiny silvery reflective surface, like that on the sides of some of the *Zaprionus* species. This distinctive characteristic allows clumping into three groups of species, of which at least one (the *sulfurigaster* clade, Wilson et al. 1969, Kitagawa et al. 1982) appears to be monophyletic. However, perhaps the best way to categorize to species or subspecies level is by their sexual display behavior. The behavior of many species was described by Spieth (1969). A more extensive comparative analysis of all recognized lineages, was made in Waddell (1990), including stocks of Taxon F, I and J, three lineages not yet formally described, but collected by Osamu Kitagawa and colleagues (Kitagawa et al. 1982).

There were two major efforts bringing stocks of these species into long-term and accessible culture. The first were those of Wilson and colleagues in the late 60’s (Wilson et al. 1969). Many of these stocks have been available via the US Stock Centers since that time. The other was 1979 and 1981 via Kitagawa and colleagues (Kitagawa et al. 1982). Some of these stocks persist in the Ehime collection and in individual laboratory collections such as Tokyo Metropolitan University. Altogether, in the 1980’s, hundreds of stocks of diverse providence were in culture, covering all known taxa resulting in dozens if not hundreds of research papers.

Active curation of these stocks—as often happens for less studied species—waxed and waned over the subsequent decades. Currently this species group is receiving renewed attention and ideally new isolates will be collected from the field. However, as much of the current work hinges on the extant stocks we believe that it is imperative that the current state of these stocks be assessed and curated.

### Incongruent phenotypes among extant stocks

As part of a follow up to Waddell (1990), we received and cultured 23 stocks (all nominally of the *nasuta*-subgroup except for 5 outgroup stocks) from the University of California San Diego Stock Center in 2013 and 2014. Over the course of receiving and establishing these stocks, we noticed several concerns. For example, a stock labeled *D. kohkoa* (15112-1771.04, Rizal, Luzon, Philippines, 1972) did not fit either the morphological or behavioral repertoire expected. It’s behavior conformed to that of *D. sulfurigaster albostrigata* (Spieth 1969, Waddell 1990), which is practically inseparable from that of *D. s. neonasuta* (Waddell 1990). Like Suzuki and Kitagawa (1990), we view *neonasuta* as a sub-population of *albostrigata*, and hence also a synonym. Further, rather than pollinosity across the whole frons, it had wide stripes of pollinosity on the frons along the orbits, exactly the morphology expected of *D. s. albostrigata*.

Another UCSD stock that was anomalous was a stock labeled *D. s. albostrigata* 15112-1811.06 from Luzon, Philippines. Unlike *nasuta* subgroup flies, which are mid-sized *Drosophila* with a honey brown color predominating, these flies were much larger with darker blackish grey tones. The behavior conformed to that of *D. siamana* of the *hypocausta* subgroup (Asada et al. 1992).

Via the Ehime stock collection and Dr Mayoshi Watada, five stocks were obtained of which three appeared anomalous. The first was labeled *D. niveifrons* from Lae in Papua New Guinea (O-30, collected 1979). This showed sexual dances specific to the *D. s. sulfurigaster* and *D. s. bilimbata* lineages, while both effectively show the same behavior (Waddell 1990). While *bilimbata* is described as a sub-species, it might well be a human spread population, or series of populations, of *D. s. sulfurigaster*. The case for its synonymy, however, is not as clear cut as with *D. s. neonasuta*. The banding patterns on the frons of this stock was of the *D. sulfurigaster* type; quite unlike that of *D. niveifrons* which has silvery pollinosity across the frons.

The stock labeled *D. pallidifrons* (PNI-75, Ponape, 1979) from Ehime did not show any pollinosity on the frons as expected. However it did not show any of the distinct sexual behaviors of the *pallidifrons* taxa (*pallidifrons*, Taxon I and Taxon J) either. It’s behavior was erratic, and infrequent, but what was observed was consistent with Taxon F, the only other *nasuta*-subgroup lineage meeting the general morphological description. This *pallidifrons* stock, as supplied by Osamu Kitagawa, is listed in Waddell (1990) as being collected June 27, 1981, and Suzuki and Kitagawa (1990) indicate that all stocks of *pallidifrons* they analyzed were from 1981. While taxon F behaved and looked as expected (Waddell 1990). The stock’s name is listed as B-208 andTaxon F was only ever recorded by Kitagawa from Borneo. Indeed, he had communicated the stock in the 1980’s with this designation (along with another stock called B-223), and both with collection details Kuching, Sarawak (Borneo), Malaysia 6/6/1971.

Finally, it seems there was only one stock of Taxon J in captivity by the late 1980’s and it was labeled Nou-98, collected August 4, 1981 in Noumea, New Caledonia. It’s behavior is described in Waddell (1990). The stock labeled taxon J Nou-98 that arrived via the Ehime center, had pollinosity across the whole frons and showed the typical sexual behavior of *D. albomicans* and *D. nasuta*. The behavior of these last two species is not visibly different, nor do they show any evidence of assortative mating (e.g., Kim et al. 2013). The main distinguishing innate feature is that *albomicans* has fused the sex chromosomes to the large autosomal chromosome three. In addition, these species are allopatric except for a hybrid zone in the region of North East India.

### Estimates of the species tree using whole genome sequencing sheds further light

We are also using data on these taxa is to develop robust and improved methods of assembly free and alignment free (AFAF) methods of phylogenetic analysis. These methods can help deconvolute problems of stock identification among cryptic species. The raw reads for these genomes are particularly well-suited to exploring AFAF analysis as they were all sequenced with the exact same chemistry, the same technology, on the same machines with mixed libraries and all have moderate to deep coverage. AFAF analyses have a number of advantages over assembly and alignment, including circumventing a range of ascertainment and other biases that can be difficult and/or time consuming to uncover.

We calculated a Neighbor-Joining tree using PAUP* (Swofford 2000) based on Poisson corrected Jaccard index-based distances. These distances were based on a random sample of 100,000 2lmers found in the raw reads using MASH (Ondov et al. 2016). A minimum kmer frequency filter of 5 was used to screen out sequencing errors and low level contamination. Kmers of length 21 have a very low probability of occurring by chance (on average, well less than one in a million in these genomes).

### Phylogeny corroboration with another approach

We validate this genome-wide species tree estimate with a novel character-based analysis we have developed. First, randomly choose a 21 mer from all those present in all sequences of all stocks. It’s presence (character C) or absence (A) is assessed across all stocks. To be kept that kmer must have a minimum frequency of 5 in at least one stock and maximum frequency of no more than twice the average coverage of that stock, across all stocks. Else, resample another kmer randomly. This was repeated until 2 million such kmers were kept. This matrix was then analyzed in PAUP* using SYD quartets (Chifman and Kubatko 2014). David Swofford has kindly extended the implementation of SYD quartets in PAUP* to deal with purine/pyrimidine DNA sequences, but here it works equally well for kmer presence/absence encoded in this way. The resulting tree with the results of 100 bootstrap replicates is shown in figure 1.

**Figure 1.**
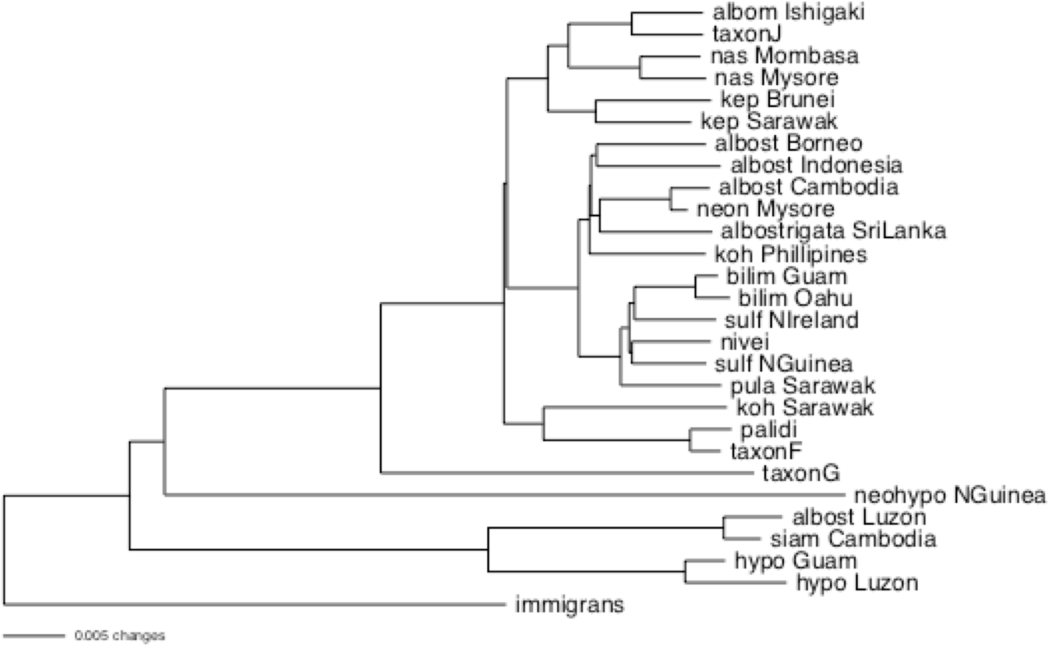
A Neighbor-Joining tree based on kmer distances.

This tree is mostly consistent with expectations based on the extensive phylogenetic analyses of Waddell (1990) analyzing a wide range of data and trees (see also the analyses of Kitagawa 1990). Note, later analyses, such as Yu et al. (1999) and Bachtrog (2006) appear compromised by mislabeled stocks and/or apparent mtDNA interspecies transfers (Waddell et al., unpublished). The clade of *pulaua* + *sulfurigaster* + *bilimbata* seen in figure 1 is consistent with some of the most parsimonious trees based on behavioral attributes (Waddell 1990). However, the very close association of *pallidifrons* with taxon F is not consistent. Our best estimate based on the phylogenetic results in Waddell (1990) and Kitagawa (1990) is that, on average, the *pallidifrons* group branches deeper than the *kohkoa* + taxon F association, but not as deep as Taxon G (a true *D. niveifrons* stock and its old name). "On average" is an appropriate qualifier, as this subgroup/species complex is primed for between species introgressions (Waddell 1990).

These trees confirm that morphological and behavioral clues reliably predict the true identity of these stocks. For *kohkoa* Philippines, the stock is clearly of the *albostrigata* lineage. Further, its location on our tree’s, which have multiple stocks from across the range of *albostrigata*, is biogeographically consistent with a stock of *albostrigata* from the Philippines. Further, as behavior and body form suggest, *albostrigata* Luzon is not of the *nasuta* subgroup and appears *primafacie* to be a stock of *siamana.* Whether it is consistent with a stock of *siamana* from the Philippines awaits a biogeographic analysis of *siamana* stocks.

Referring to our species trees and the case of the *D. pallidifrons* stock PNI-75 (not PNl 75 as some have written), the evidence, particularly the low level of genetic divergence, supports the hypothesis that this is a wrongly labeled stock of Taxon F. In the case of the Taxon J stock, it indeed appears to be a stock of *D. albomicans.* That the mtDNA of this stock reported in Yu et al. (1999) clusters with *D. albomicans* sequences suggests that the confusion of this stock may be quite early, but not as early as 1988 when this stock was observed to show a distinct behavior and morphology quite like true *D. pallidifrons* (Waddell 1990). In the case of the stock labeled *D. niveifrons*, this indeed appears to be a stock of *D.* s. *sulfurigaster* or *D.* s. *bilimbata.* That this stock often clusters sister to a *D.* s. *sulfurigaster* stock from Wau in Papua New Guinea, suggests that the recorded collection locality of this mislabeled stock might indeed be correctly retained, that is, Lae, Papua New Guinea.

Concerns over the *D. albostrigata* Cambodia stock (15112-1811.04, Siam Reap, Cambodia, 1968) show up in figures 1 and 2. Based on biogeography and the pattern of shared chromosomal inversions (Suzuki and Kitagawa 1990), *neonasuta* should be just another stock of *albostrigata* from southern India. That being the case, it is expected to be most closely related to Sri Lankan *albostrigata.* This is not the case and the Cambodian stock is of surprisingly low genetic differentiation from *neonasuta* Mysore; an anomaly worth investigating.

**Figure 2.**
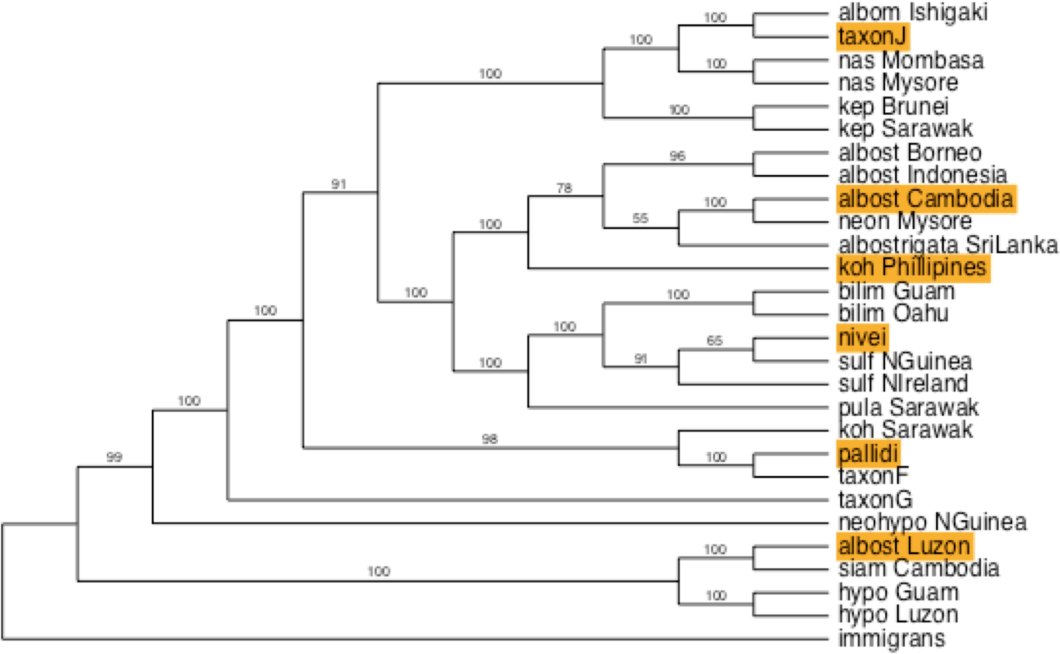
The SVD quartets-based tree from PAUP* based on 2 million randomly sampled 21-mers from across the genomes of these stocks. Support values are based on 100 bootstrap resamplings of the data matrix. Highlighted are the stocks we diagnose as mislabeled; in the case of *albostrigata* Cambodia, we suspect stock introgression.

### Diagnosing the anomalous *albostrigata* Cambodia

In a situation like this, a look at the NeighborNet (Bryant and Moulton 2004) representation can be very useful. NeighborNet measures discrepancies of the evolutionary distances between taxa to detect non-tree signals, such as those due to introgression or lineage mixing. It then aims to show these discrepancies in the form of a planner graph (a diagram, that can be written on a page without need of 3 dimensions to represent all predicted distances). For an explanation of how NeighborNet results relate to other tests of possible introgression, such as ABBA-BABA tests, see Waddell (2018). That article also describes how the genomic sequence of a single non-admixed individual can well represent its ancestral population in this type of analysis, due to its genome being many effectively unlinked genetic loci from just that population.

The NeighborNet of just the *sulfurigaster* group stocks (figure 3) seems a very good representation, showing over 99.975% of the variance in the data. It is clear that the largest non-tree splits involve *albostrigata* Cambodia sharing genetic material similar to that of *albostrigata* Indonesia and *albostrigata* Borneo. Consistent with this view, when this stock is removed from the analysis, the NeighborNet becomes much more tree-like (figure 3). Now, this *neonasuta* stock is segregated much more strongly with *albostrigata* SriLanka. This does not prove that there was stock contamination, for example, but is highly suggestive that either this occurred or the populations of *albostrigata* in Cambodia and/or southern India have a complex evolutionary history somewhat in contradiction to the biogeographic expectations and the structure seen with other sampled populations of *albostrigata.* Note, there are other non-tree signals also showing up, with the most prominent and biogeographically understandable being *pulaua* from Borneo apparently sharing more alleles with some *albostrigata* stocks than with others. These species are sympatric, interfertile and will mate with each other in confinement (Kitagawa et al. 1982), introducing the possibility of interspecies introgression in the wild.

**Figure 3.**
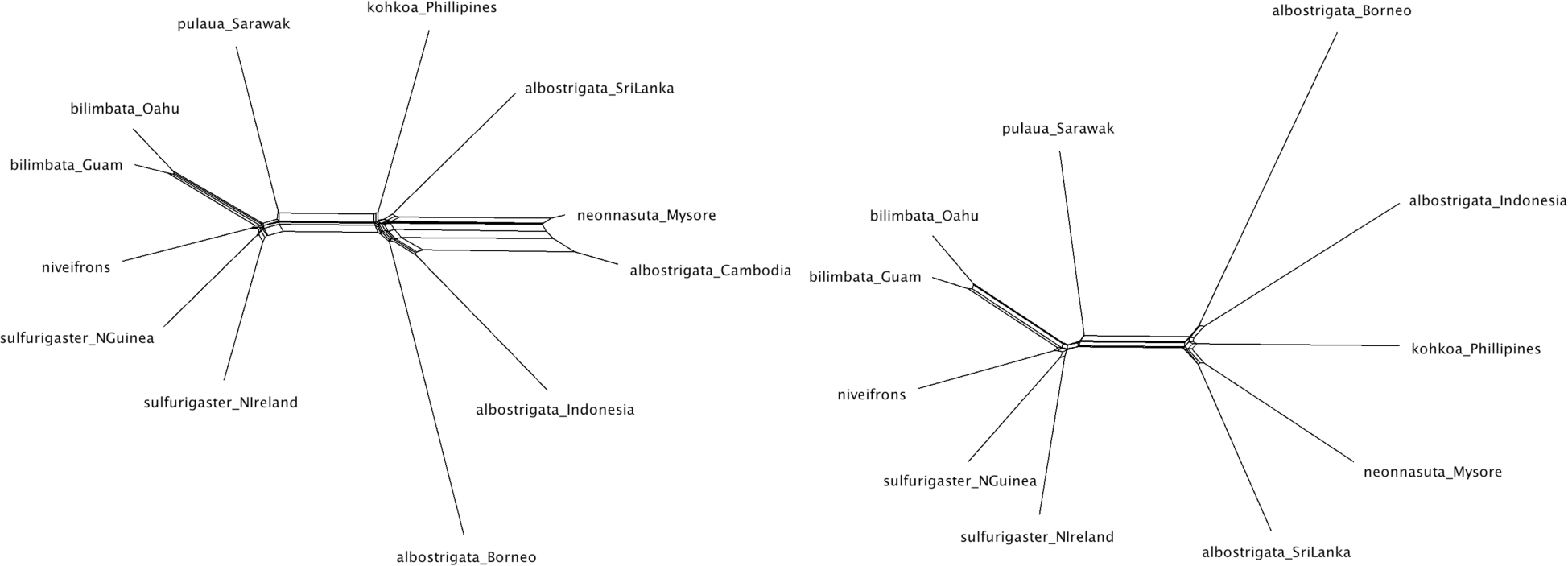
(Left) The NeighborNet, via SplitsTree4, of *sulfurigaster* group stocks based on whole genome 2 lmer Poisson corrected distances passing a minimum 5 and maximum 100 filter. The fraction of the total variance of the data explained by the graph is 0.99975. (Right) With *albostrigata* Cambodia removed. The fraction of the total variance ofthe data explained is 0.99973.

## Conclusions

The diverse natural history, unique phenotypes, sexual dynamism and volatile chromosomal biology of the *nasuta* group all bring interest to the group making it useful to have stocks of this group available for study. At the same time, long term culture runs the risk of mislabelling of stocks and/or introgressing stocks. In this sample of 28 stocks, detected rates of such problems are as high as 6 out of 28 or ~21%. These issues were reported to stock centers at the time, but due to limited resources and declining Federal support little could be done. Since then both stock centers are now under new management, and in new locations. The US Drosophila Species Stock Center has discontinued nearly all their *nasuta-subgroup* stocks, while many of the original Ehime stocks continue at the Kyorin-fly stock center in Japan.

We suspect that the geographic locations of stocks that often appears as a second part of the stock description has been both a hinderance and a help. The first because, if it is the only recognized part of a label, it could lead to the assumption it is another species sharing at least part of the locality name, something that may have happened with some of these mislabeled stocks. A help, because in many cases, along with reliable genetic data, it may lead to a dominant probable hypothesis of what the stock really is.

Finally, we call on the larger *Drosophila* community to make a push for continued robust support of our stock centers, to reassess and curate extant stocks, and to invest in genetic barcoding of stocks when they are accepted into long-term culture. Mislabelling and related issues such as these can have lasting impact. For instance, these mixed up stocks may have compromised the results of a number of papers. For example, in the paper of Yu et al. (1999) the taxon J stock shows an *albomicans* type mtDNA sequence further confusing an already confused mtDNA tree (due to multiple real introgressions, Waddell et al. unpublished). The stock marked *D. kohkoa* Rizall Philippines that we have diagnosed as a mislabeled *albostrigata* stock was also used inin a variety of papers..

When authors lodge their sequences in GenBank this allows these possibilities, along with the other conclusions of their papers to be tested. In the case of Yu et al. (1999), this was done so we can check and we observe that the mtDNA sequence of their Taxon I and J is identical to our Taxon J, while their *pallidifrons* shows two transversion differences suggestive of sequencing error. All cluster tightly on trees to the exclusion of all other stocks, suggesting all three are mislabeled *albomicans*, possibly the very same stock. *That said, both the high rates of stock mislabelling and the failure to spot such errors in peer reviewed publications reinforces the need for good biological practices of species identification before reporting DNA sequences or analyses*. Further, flagging anomalous stocks such a *D. s. albostrigata* Cambodia requires the aforementioned background of properly identified stocks along with diverse genetic data and an analysis sensitive enough to detect possible genetic mixing. Both types of diagnosis are essential if either phylogenetic or genetic conclusions from long cultured stock studies are to be trusted.

## Acknowledgements

Sincere thanks to Prof. Osamu Kitagawa for the wide range of stocks he made available and his extensive efforts into understanding this exceptional subgroup of *Drosophila*. To Mayoshi Watada (Ehime) and the UCSD Stock Center for sending stocks most recently. This work was partly supported by NIH grant 5R01LM008626 to PJW.

